# Mycolactone A vs. B: Does localization or association explain isomer-specific toxicity?

**DOI:** 10.1101/2023.05.19.541532

**Authors:** John D. M. Nguyen, Gabriel C. A. da Hora, Jessica M. J. Swanson

## Abstract

Mycolactone is an exotoxin produced by *Mycobacterium ulcerans* that causes the neglected tropical skin disease Buruli ulcer. This toxin inhibits the Sec61 translocon in the endoplasmic reticulum (ER), preventing the host cell from producing many secretory and transmembrane proteins, resulting in cytotoxic and immunomodulatory effects. Interestingly, only one of the two dominant isoforms of mycolactone is cytotoxic. Here, we investigate the origin of this specificity by performing extensive molecular dynamics (MD) simulations with enhanced free energy sampling to query the association trends of the two isoforms with both the Sec61 translocon and the ER membrane, which serves as a toxin reservoir prior to association. Our results suggest that mycolactone B (the cytotoxic isoform) has a stronger association with the ER membrane than mycolactone A due to more favorable interactions with membrane lipids and water molecules. This could increase the reservoir of toxin proximal to the Sec61 translocon. Isomer B also interacts more closely with the lumenal and lateral gates of the translocon, the dynamics of which are essential for protein translocation. These interactions induce a more closed conformation, which has been suggested to block signal peptide insertion and subsequent protein translocation. Collectively, these findings suggest that isomer B’s unique cytotoxicity is a consequence of both increased localization to the ER membrane and channel-locking association with the Sec61 translocon, facets that could be targeted in the development of Buruli Ulcer diagnostics and Sec61-targeted therapeutics.

## INTRODUCTION

Buruli ulcer is a neglected tropical disease caused by *Mycobacterium ulcerans* that leads to large necrotic skin ulcerations that surprisingly lack pain and inflammation (1, 2, 3). These pathogenic effects are caused by a macrolide exotoxin produced by the bacteria known as mycolactone. The toxin is believed to be secreted in bacterial vesicles (4) and delivered to host cells where it induces cytotoxic and immunosuppressive effects by disrupting several cellular processes such as cytoskeletal organization, protein production and signaling cascades (2, 4, 5). Structurally, mycolactone consists of a 12-membered lactone ring with two highly unsaturated chains (**Figure 1**). The southern chain has been shown to play a key role in the toxin’s cytotoxicity; first through mycolactone variants that differ in bioactivity possessing the same northern chain and lactone ring structure but varying in the southern chain (2, 6, 7). Additionally, it has been reported that synthetic mycolactone subunits lacking the northern chain partially retained immunosuppressive activity, while subunits lacking the southern chain or both chains were inactive (8). Although these structure-activity relationships have been reported, the underlying mechanisms remain unknown.

**Figure 1.**
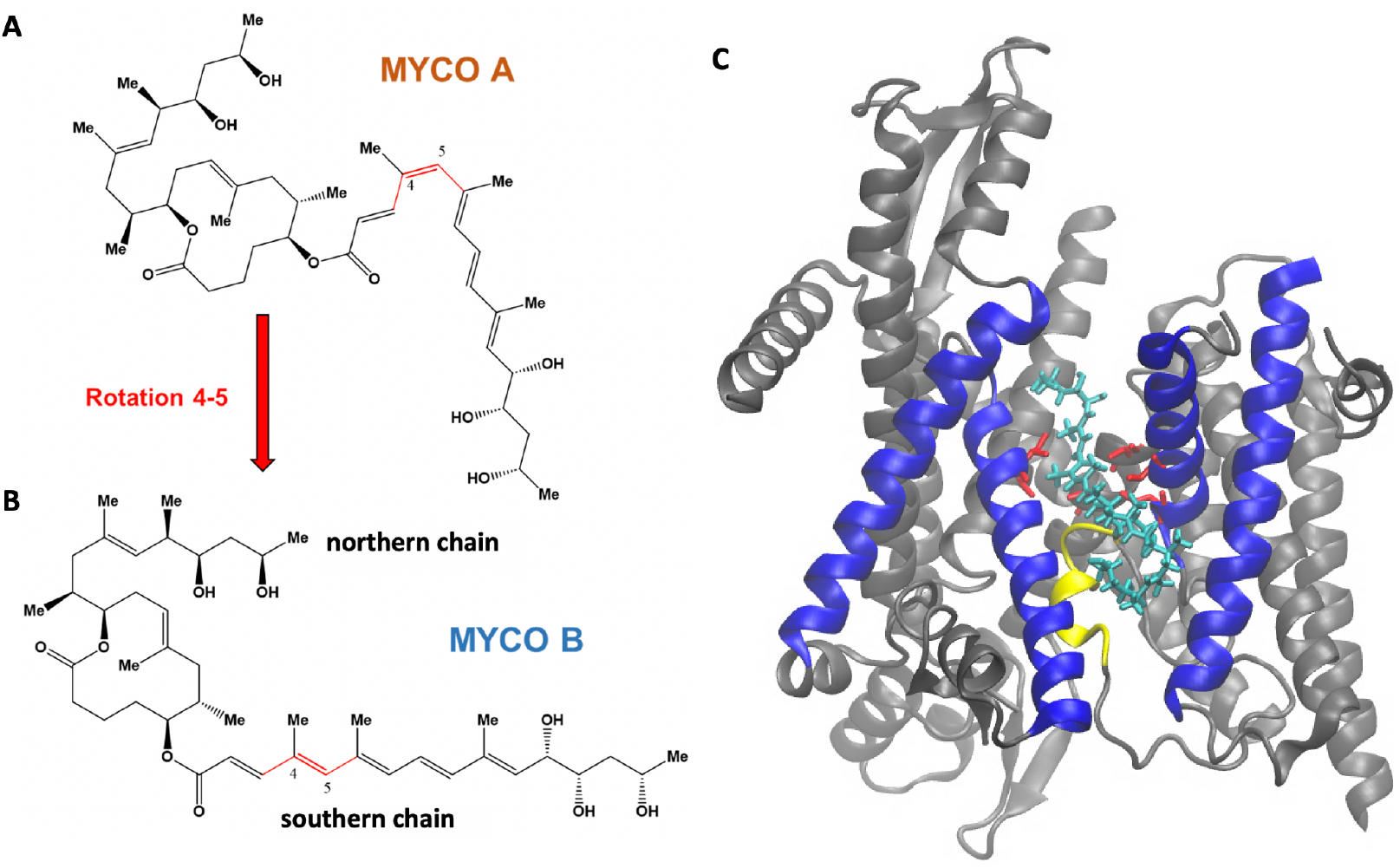
Structures of mycolactone A **(A)** and B **(B)** identifying the northern and southern chains and dihedral angle 3-4-5-6 (cis and trans, respectively) in red. **(C)** The Sec61 translocon with mycolactone B (cyan) bound, highlighting the plug (yellow), lateral gate (blue: TMs 2b, 3, 7 and 8) and pore ring residues (red).

Mycolactone exists primarily as two cis/trans stereoisomers, designated as mycolactone A and mycolactone B, respectively, which are in dynamic equilibrium with a 60:40 ratio (9). These two isoforms differ in a rotation around the toxin’s 4-5 double bond in the southern chain (**Figure 1**). The bioactivity of the two isomers was studied by synthesizing analogs that closely resemble the natural compounds but with slight structural variations, while preserving the cis/trans aspect of the southern chain. Results showed that the B analog was significantly more cytotoxic, suggesting that mycolactone B is the primary virulence factor of Buruli ulcer (10). Interesting possible explanations of this isomeric specificity include differences in adsorption, cellular localization and/or host target interactions.

Mycolactone was initially believed to enter the host cytosol following passive diffusion through the cell membrane (11). However, computational studies initially predicted (12), and experimental work showed that the toxin strongly interacts with cellular membranes (13). Nitenburg et al. (13) studied mycolactone’s interaction with membranes using Langmuir monolayers and reported that the toxin interacts with membranes at very low concentrations and disturbs lipid organization. Our previous work quantified a strong free energy of association between mycolactone and phospholipid membranes and revealed the toxin’s preference for the interfacial region of bilayers below the lipid headgroups (12, 14). Most recently, we demonstrated that mycolactone B has a stronger association free energy with the ER membrane than with a model plasma membrane (15). Moreover, the toxin reorganizes the plasma membrane to reflect the more disordered ER-like composition locally. Given recent work demonstrating the role of lipid order trafficking proteins (16), it is possible that the toxin’s preferential lipid interactions influence its cellular localization. Given these findings, it is likely that what was previously believed to be the toxin’s localization in the cytosol was more likely the association of mycolactone with the ER membrane.

Mycolactone acts by invading host cells and interacting with multiple intracellular targets, including the angiotensin II receptor, the Wiskott-Aldrich syndrome protein, mTOR, and the Sec61 translocon (17, 18, 19, 20). Of these identified targets, the most significant is the Sec61 translocon, as mycolactone’s cytotoxic and immunosuppressive effects primarily result from its inhibition (6). The Sec61 translocon is a membrane-embedded protein complex (**Figure 1**) that translocates precursor polypeptides into the ER for processing. The toxin binds to the pore-forming Sec61α subunit of the protein and strongly inhibits its ability to translocate polypeptides. This reduces the cell’s ability to produce many secretory and transmembrane proteins, which leads to multiple downstream consequences, such as immunomodulation and cytotoxic effects (1, 21). Single amino acid mutations in Sec61 have been identified that confer cells broad resistance to mycolactone’s cytotoxic and immunomodulatory effects while not affecting the protein’s functionality (6).

A recent study by Itskanov et al. (22) reported near-atomic-resolution cryo-electron microscopy (cryo-EM) structures of the human Sec61 translocon bound by seven small molecule inhibitors – including mycolactone. Although each inhibitor in the study possessed distinct chemical structures, they all interact with hydrophobic side chains of three key gating elements of Sec61 referred to as the plug, lateral gate, and pore ring (**Figure 1**). The plug is a short helix that must be displaced to allow translocating peptides access to the ER lumen, while the lateral gate consists of transmembrane segments (TMs) 2b, 3, 7, and 8 of the channel that can separate and open the channel laterally to allow peptides access to the ER membrane environment (23). Each of the inhibitors studied by Itskanov et al. (22) interacts with TMs 2b and 3, which form one side of the lateral gate, as well as TM 7, forming the other side. Given that concerted movement of the plug and lateral gate is required for the insertion of an incoming signal sequence and subsequent translocation, this coordination of interactions likely blocks signal insertion. The pore ring is a constriction of six hydrophobic residues (I81, V85, I179, I183, I292, and I449) that translocating peptides pass through and is gated by movement of the plug (23). In addition to hydrophobic interactions with the three gating elements of Sec61, the authors also found that all of the inhibitors form polar interactions with a polar cluster of residues of the lateral gate, which was confirmed by mutational analysis of yeast equivalents of these residues to be key for binding affinity (22). Taken together, Itskanov et al.’s (22) findings show that all seven tested Sec61 inhibitors act by locking the lateral and lumenal gates of the channel to prevent dynamics required for translocation.

In this work, we query the molecular mechanisms underlying mycolactone’s bioactivity by comparing the interactions of the two isomers with a model ER membrane and with the Sec61 translocon. We use MD combined with Transition-Tempered Metadynamics (TTMetaD) enhanced sampling in order to calculate the permeation free energy profiles and characterize the association of mycolactone A and B with a model ER membrane, aiming to investigate the role of localization in the toxin’s pathogenicity. The ER membrane is chosen since it is the likely reservoir for the toxin prior to its association with the Sec61 translocon. We also use MD to model mycolactone A/B-Sec61 complexes embedded in an ER membrane to identify the existence of interactions and/or dynamics that may contribute to isomeric specificity. We first present permeation free energy profiles, which show a stronger affinity of mycolactone B for the ER membrane than mycolactone A. This has interesting ramifications for both cellular localization and competition between the isomers for the translocon. We find that isomer B has a more open structure that allows it to better interact with the membrane lipids and water molecules while in the membrane compared to the A isomer’s more compact structure. Comparing simulations of the toxin-Sec61 complexes, we find that mycolactone A is unable to interact with the plug domain and TMs 7 to the extent that mycolactone B does – suggesting isomer B’s increased ability to lock the plug domain and TMs 2, 3, and 7 in place to inhibit translocation. We also find that mycolactone B has more interactions with the pore ring and is more inserted into the constriction, which may result in a stronger binding affinity and increased ability to occlude the pore from peptides. Additionally, polar interactions are better satisfied with mycolactone B bound via interactions with the polar cluster residues of the lateral gate and water molecules. Lastly, the isomer B-bound complex adopts a more closed conformation on the cytosolic side of the translocon than the A isomer, which may be linked to mycolactone B’s increased cytotoxicity.

## RESULTS AND DISCUSSION

### Endoplasmic reticulum membrane association

Two-dimensional free energy profiles (potentials mean of force; 2D-PMFs) for the permeation of mycolactone A and mycolactone B through the ER membrane are shown in **Figure 2**. The two collective variables (CVs) that were tracked are the permeation depth, tracked via the z-distance between the center of mass (COM) of the lactone ring and that of the membrane, and the orientation, tracked via the angle between the vector connecting the northern and southern chains’ hydroxyl groups and the membrane normal (15). The black lines track the minimum free energy paths, which are the most probable paths taken by the isomers through the membrane. Inserted snapshots (**Figure 2D,E**) show the dominant configurations in minima along the minimum free energy paths. In these minima, the polar hydroxyl groups at the ends of each chain hydrogen bond with oxygen atoms from lipid headgroups, glycerol and/or water, while the macrolide ring and aliphatic portion of the southern chain interact with the hydrophobic lipid tails. This is consistent with our previous computational work, where we studied mycolactone B’s interaction with other model membranes (12, 14, 15).

**Figure 2.**
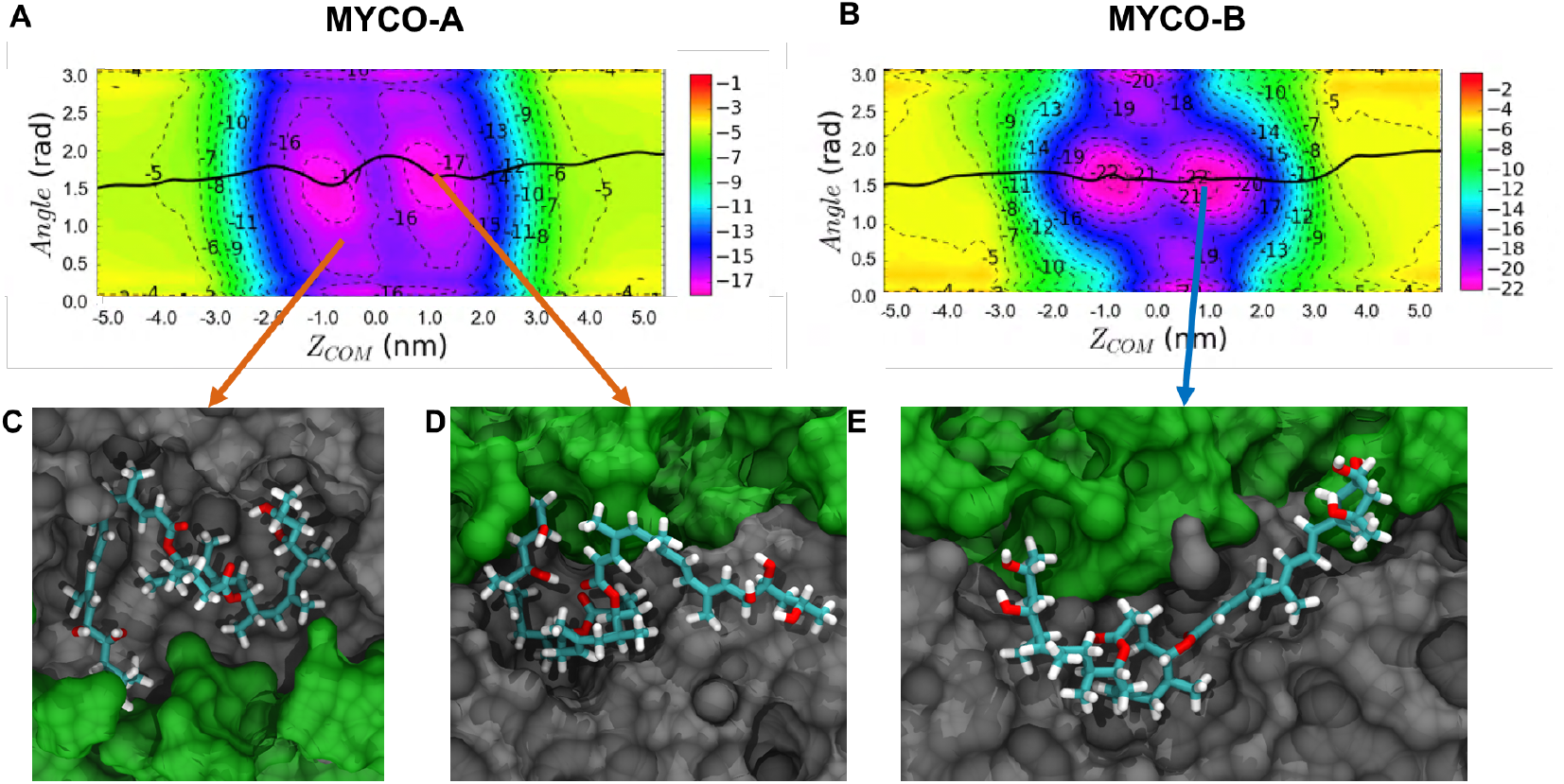
2D free energy profiles of the permeation of **(A)** mycolactone A and **(B)** mycolactone B through the ER membrane. The black lines trace the toxin’s most probable paths. **C-D**) representative configurations. The toxin is colored by atom type, lipid headgroups, and tails are colored green and gray, respectively. Water is omitted for clarity. The membrane spans -2.0 < *Z*_*COM*_ < 2.0 nm. The energy is shown in kilocalories per mole (kcal/mol).

The minimum free energy paths of both isomers through the ER membrane are similar: the toxin first adopts conformations that optimize polar and nonpolar interactions at the interfacial region of the membrane below the lipid headgroups, and then migrates from one leaflet to the other by swapping its polar interactions with lipid headgroups of one layer to lipid headgroups of the opposite leaflet. However, while associating with the membrane, it appears that isomer A’s compact structure has fewer favorable interactions with the membrane and more intramolecular interactions (**Figure 2D**), while isomer B’s more open structure demands more interactions with the bilayer and/or water (**Figure 2E**). There is also a difference in the binding affinity, with mycolactone B having a more favorable interaction with the ER membrane than mycolactone A. This is more evident in the corresponding one-dimensional free energy profiles (**Figure 3**). Additionally, there is a difference in the membrane-associated conformational ensemble; mycolactone A has a broader angle CV distribution than mycolactone B (**Figure 2**). The free energy landscapes in **Figure 2A,B** were validated by analyzing the distribution of the CVs in unbiased simulations and found to be consistent with mycolactone A exhibiting a broader angle distribution (**Figure S1**). The extended regions in the angular distribution correspond to configurations where only one of the hydroxyl-containing chains is interacting with the lipid head groups or glycerol, while the other is buried in the lipid tails, satisfying polar groups with intramolecular interactions (**Figure 2D**). This observation suggests that the interactions between mycolactone A’s hydroxyl tail ends and the lipid headgroups are more transient, whereas mycolactone B’s are more consistent.

**Figure 3.**
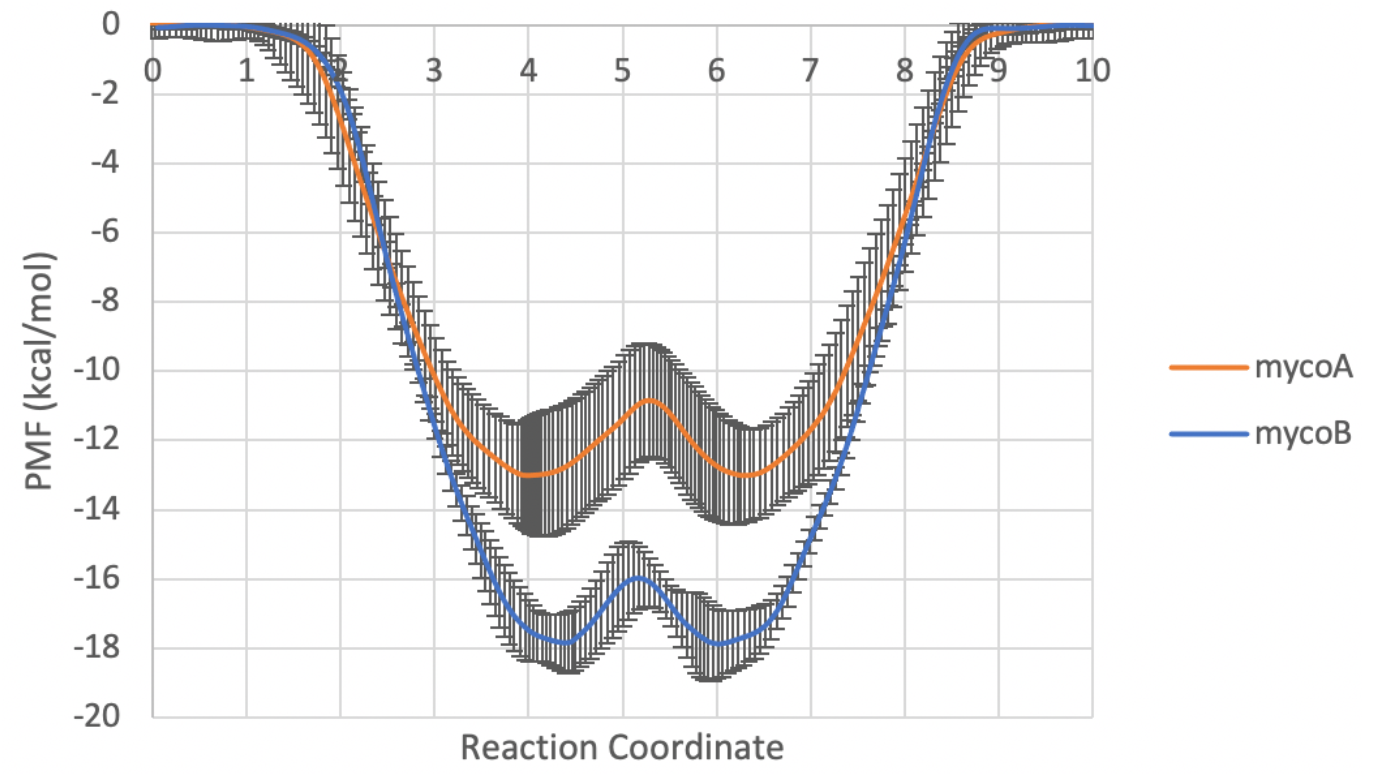
1D free energy profiles of each system along the minimum free energy path. The standard errors range from 0.03 to 1.82 kcal mol^-1^ for mycolactone A and from 0.06 to 1.22 kcal mol^-1^ for mycolactone B.

To further query how the observed difference in binding affinities arises from the structural differences between the isomers, we turn to interaction energies. With a cis double bond in the southern chain, mycolactone A is more compact, which may shift interactions with its environment to interactions with itself. To test this, the GROMACS tool *gmx_energy* was used to calculate the average interaction potential energies between the toxin and the ER membrane and water molecules, as well as the intramolecular toxin interactions (**Table 1**). Energy values were calculated for configurations where mycolactone is in the membrane hydrophobic core (−2.0 < Z < 2.0 nm). From this analysis, it appears that while associating with the membrane, mycolactone B has a stronger interaction with the lipids and water molecules, while mycolactone A indeed has stronger intramolecular interactions.

**Table 1.**
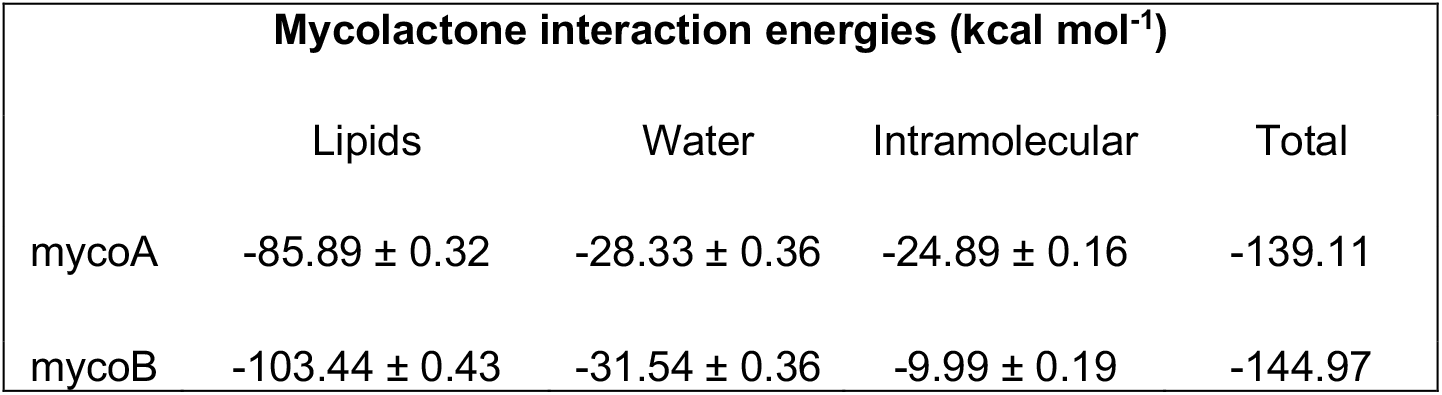
Average interaction potential energy of mycolactone with different components of the system.

| Mycolactone interaction energies (kcal mol<sup>-1</sup>) |  |  |  |  |
| --- | --- | --- | --- | --- |
|  | Lipids | Water | Intramolecular | Total |
| mycoA | -85.89 ± 0.32 | -28.33 ± 0.36 | -24.89 ± 0.16 | -139.11 |
| mycoB | -103.44 ± 0.43 | -31.54 ± 0.36 | -9.99 ± 0.19 | -144.97 |

We further evaluated the interactions between the toxin and water molecules by calculating the probability distribution of the number of water molecules in contact with the toxin as a function of the z-component distance between the lactone ring and the center of the membrane (**Figure 4**). A contact is defined as any distance between an oxygen atom of water and any oxygen atom of mycolactone less than or equal to 3.04 angstroms. Our previous work (14, 15) has shown that mycolactone B coordinates with water molecules during association with pure DPPC and model ER and plasma membranes to satisfy polar interactions as the toxin moves into hydrophobic regions of the bilayer. Here, both mycolactone A and B are able to interact with water molecules while permeating through the ER membrane, even while near the center of the membrane. However, mycolactone B is more likely to coordinate with more water molecules than mycolactone A in the ER membrane. This again is due to mycolactone A’s more compact structure, increasing its intramolecular hydrogen bonding instead of interacting with water molecules. These differences in water coordination likely contribute to mycolactone B’s stronger affinity for the ER membrane compared to isomer A.

**Figure 4.**
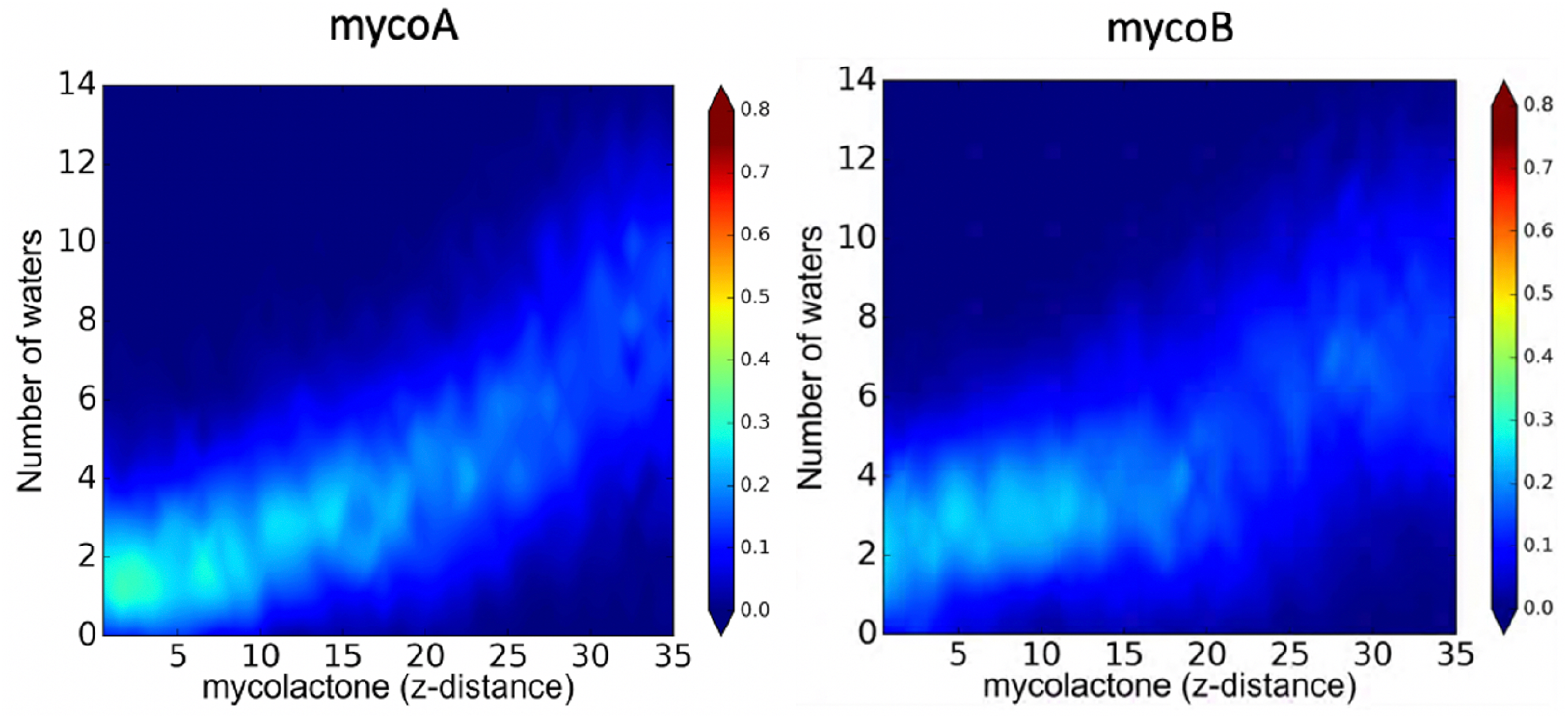
Probability distribution of the number of water molecules interacting with mycolactone with respect to the toxin position in the ER membrane in biased simulations. Z-distance is shown in angstroms.

Additionally, to investigate each isomer’s effect on the ER membrane’s physical properties, lipid tail order parameters were calculated for lipids near the toxin (within 5 angstroms) in unbiased simulations and compared to tail order parameters of an ER membrane-only simulation (**Figure S2**). Both mycolactone A and B were found to disrupt lipid tail order to a comparable degree.

### Sec61-Toxin Complex

Since the Sec61 translocon is a primary target of mycolactone and the toxin’s cytotoxic effects can mostly be attributed to this interaction, we next evaluated each isomer bound to Sec61 embedded in an ER membrane. The isomers were bound to a region of Sec61 that was identified to be mycolactone’s binding site by Itskanov et al. (22) via cryo-EM (**Figure 5**). Since only mycolactone B was modeled into the cryo-EM density, mycolactone A was docked as described in Methods. Once bound, both isomers were oriented with their southern chain inserted into the core of the translocon while the northern chain protrudes out into the hydrophobic lipid tail region of the bilayer. Each complex was simulated until the root-mean-squared deviations (RMSDs) of the protein backbone stabilized (1.2 and 1.5 microseconds for mycolactone A and B, respectively) (**Figure S3**) and the last 300 nanoseconds was used for analysis. Both isomers remained bound to the translocon throughout the simulations.

**Figure 5.**
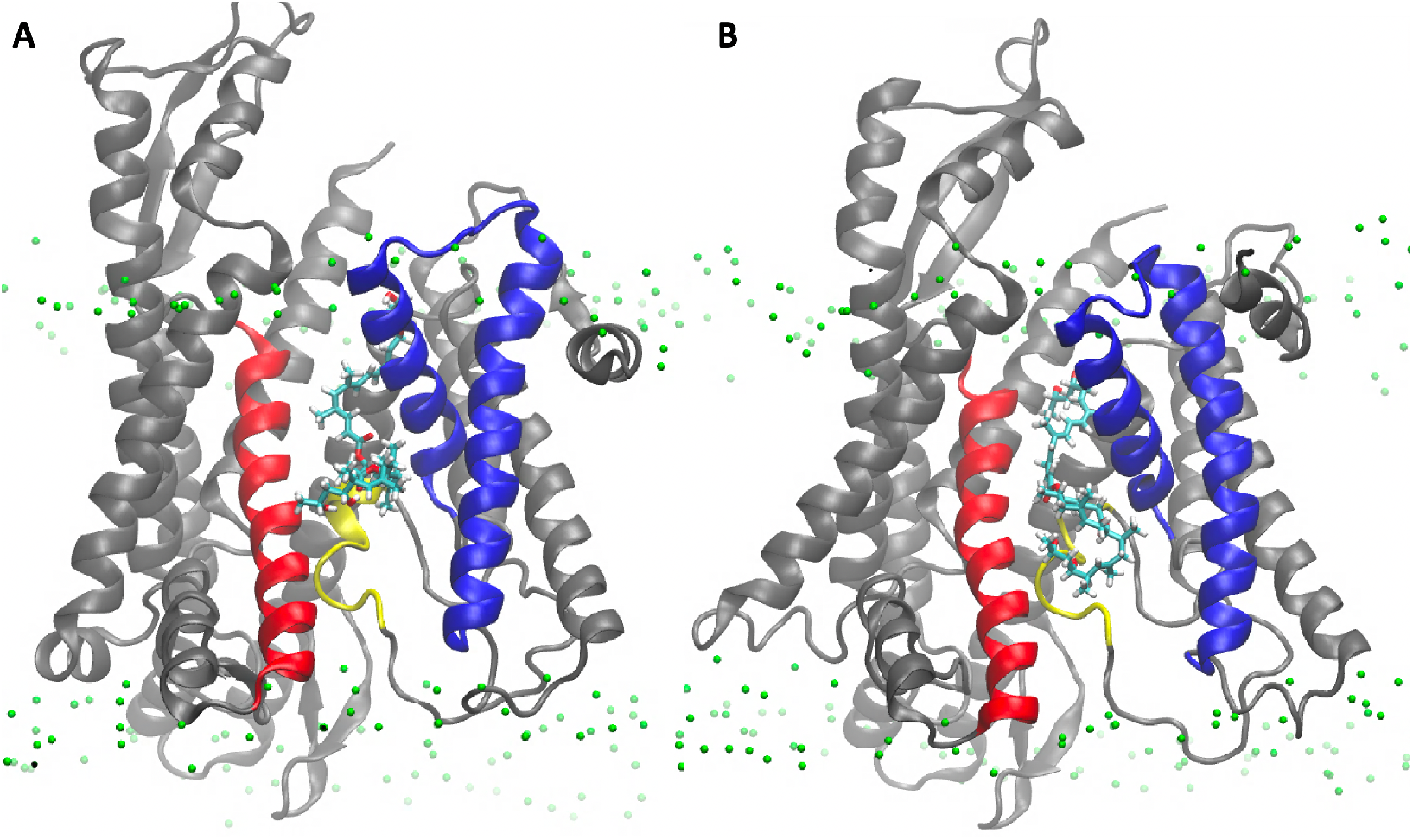
Structures of mycolactone A-Sec61α **(A)** and mycolactone B-Sec61α **(B)** complexes. Mycolactone is shown as licorice sticks and colored by atom type. Sec61α is shown as cartoon ribbons with the plug, TMs 2b and 3, and TMs 7 are colored yellow, blue, and red, respectively. Lipid head group phosphates are colored green.

We first analyzed interactions with the three key gating elements: the plug, lateral gate and pore ring. Contact analyses were performed between mycolactone isomers A/B and residues of each gating element. A contact is defined as any distance between a heavy atom of the toxin and a heavy atom of the protein less than or equal to 4.5 angstroms. Mycolactone B was found to have many contacts with the plug and both sides of the lateral gate (TMs 2b and 3 on one side and TM 7 on the other), while mycolactone A predominantly interacts with just TMs 2b and 3 (**Figures 6A-C** and **7**). The A isomer’s lack of interactions with the plug and TM 7 likely results in a weaker ability to cement the plug and two sides of the lateral gate together at their interface to restrict dynamics and inhibit peptide translocation. Mycolactone B was also found to have more contacts with the pore ring as it intercalates into the crescent-shaped constriction, whereas mycolactone A is bound further away (**Figures 6D** and **7**). This difference in interaction with the pore ring may result in a stronger binding affinity of isomer B and a possible increased ability to block the pore from peptides.

**Figure 6.**
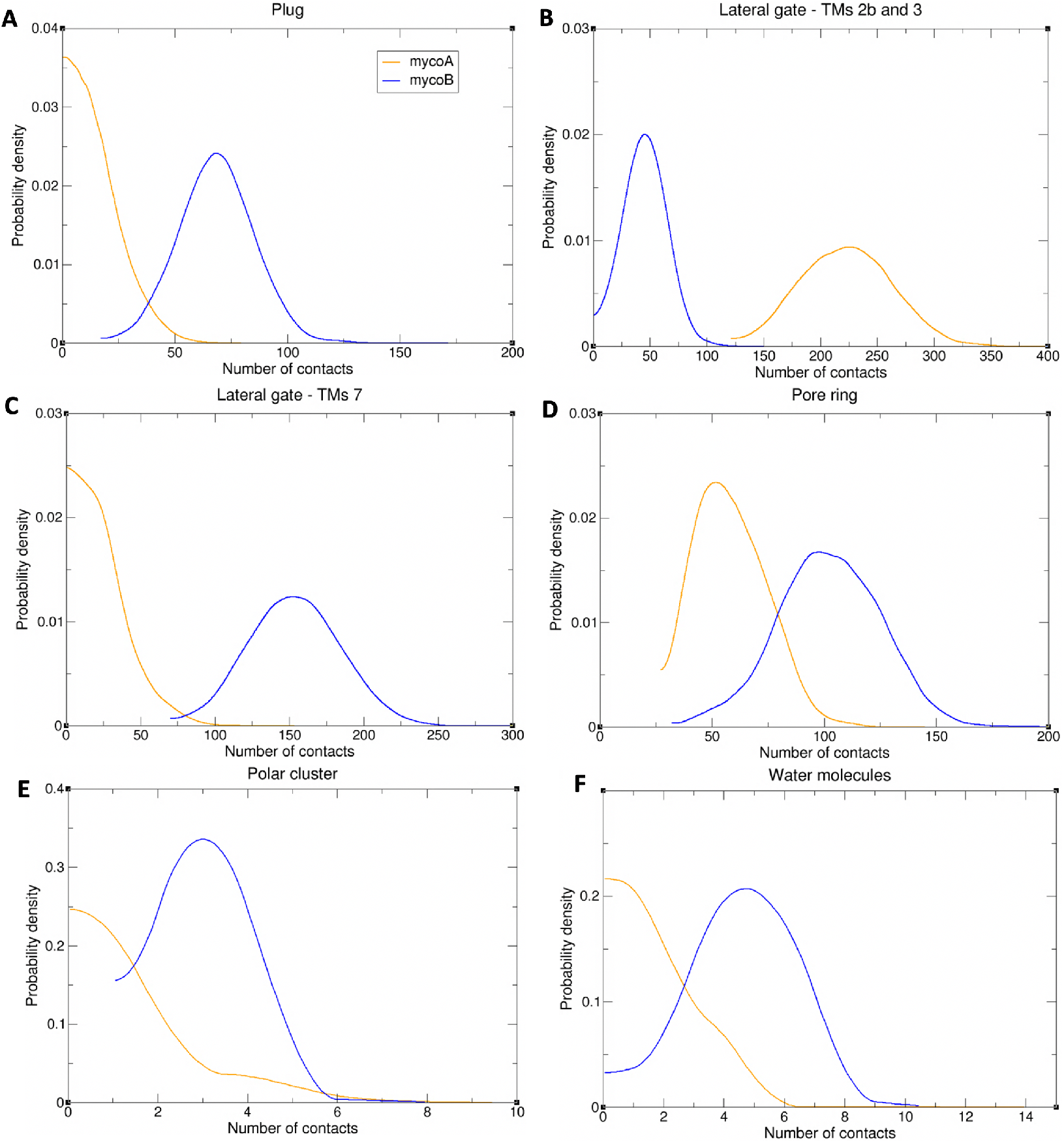
Probability of the number of contacts between mycolactone and the plug **(A)**, TMs 2b, 3 **(B)**, and 7 **(C)**, and the pore ring **(D)**, polar cluster residues **(E)**, and water molecules **(F)**.

**Figure 7.**
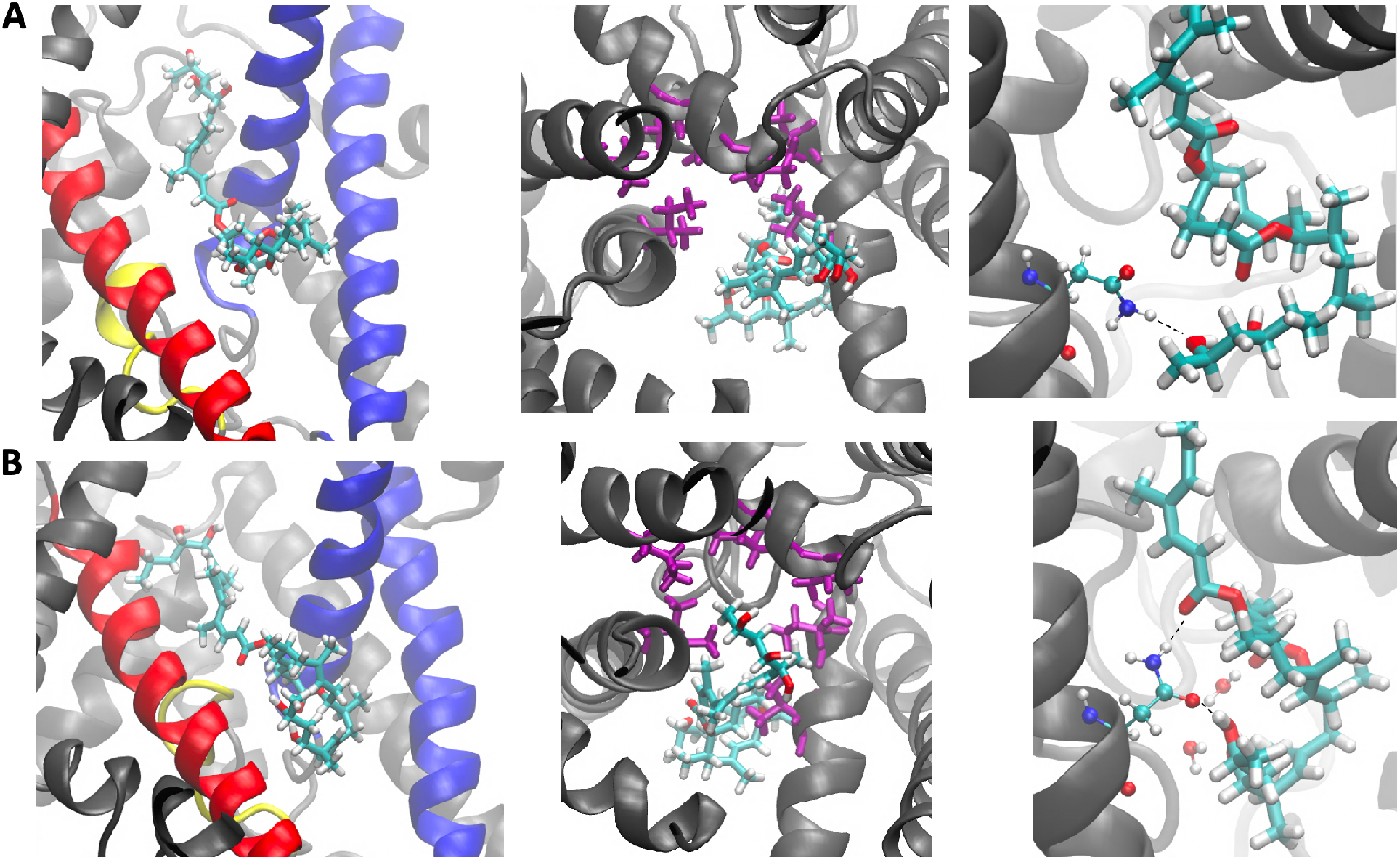
Positions of mycolactone A (**row A**) and mycolactone B (**row B**) relative to the plug, TMs 2b, 3, and 7 (left), the pore ring (middle), and N300 with coordinating water molecules (right). Mycolactone, the plug, and TMs 2b, 3, and 7 are represented the same as in Fig 5. The pore ring residues are shown as purple licorice sticks. N300 and water molecules are shown as balls and sticks and colored by atom type.

To evaluate the key polar interactions shared by all inhibitors and demonstrated as important for binding via mutational analysis by Itskanov et al. (22), the number of contacts between oxygen atoms of the toxin and oxygen and nitrogen atoms of the polar cluster residues of the lateral gate (Q127 and N300) was calculated. Mycolactone B was found to have more polar interactions with the polar cluster residues than mycolactone A, which likely contributes to a stronger binding affinity of the B isomer (**Fig 6E** and **7**). Interestingly, effectively all of the contacts calculated for both isomers belonged to interactions with N300 (100% and 99.7% for mycolactone A and B, respectively) – suggesting that N300 is more crucial for the binding affinity than Q127. This is consistent with Itskanov et al.’s (22) mutational analysis, which revealed that an N300L mutation conferred stronger resistance than a Q127L mutation to Sec61 inhibitors: cotransin, decatransin, and ipomoeassin F. In addition to polar interactions with the polar cluster residues, Itskanov et al.’s (22) mycolactone-bound structure also revealed a water molecule coordinated by the northern chain and ring of the toxin and residues of the lateral gate. This water coordination satisfies polar interactions of the northern chain and ring of the toxin that are exposed to the lipid environment. Thus, the number of contacts between oxygen atoms of the northern chain and ring of the toxin and oxygen atoms of water molecules was calculated for each isomer. Similar to interactions with the polar cluster residues, mycolactone B’s lipid exposed region was found to have more interactions with water molecules than mycolactone A (**Figures 6F** and **7**). Together, this difference in polar interactions likely contributes to a stronger binding affinity of the B isomer.

Next, we assessed the conformational changes of Sec61 induced by each isomer and found that the cytosolic side of the translocon adopts a more closed conformation with mycolactone B bound (**Figure 8**). This observation bears implications regarding the difference in toxicity. Itskanov et al.’s (22) inhibitor-bound structures of Sec61 demonstrate the conformational plasticity of the binding pocket as the width of the lateral gate opening varies depending on the bound inhibitor. Among the seven inhibitor-bound structures, the lateral gate of the cotransin-bound structure was found to have the widest opening on the cytosolic side. It has been shown that cotransin is less effective in inhibiting the translocation of peptides that have a stronger targeting signal (24, 25, 26). Itskanov et al. (22) proposed that the reason behind this observation could be the wider conformation of the lateral gate observed in their cotransin-bound structure, which might enable specific interactions between the lateral gate and a signal sequence, ultimately opening the lateral gate further and potentially causing the release of the inhibitor. Given these findings, it’s plausible that a more closed conformation of the translocon observed in our simulations is associated with mycolactone B’s increased toxicity through a similar mechanism.

**Figure 8.**
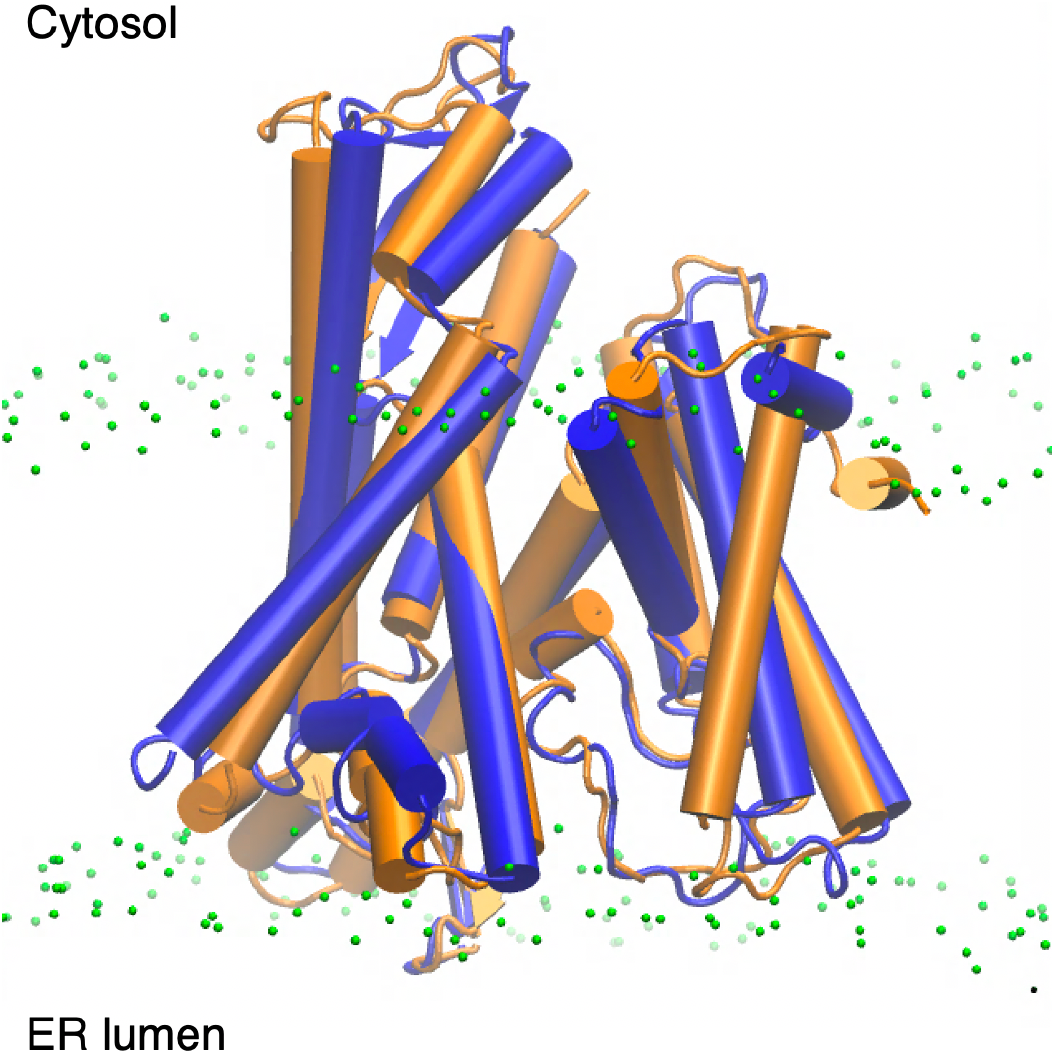
Conformations of Sec61 with mycolactone A (orange) and mycolactone B (blue) bound. Lipid head group phosphates are colored green.

## CONCLUSIONS

Using all-atom MD simulations, we have herein probed the isomeric specificity of the macrolide exotoxin mycolactone, the sole causative agent in Buruli ulcer disease. Although mycolactone is produced by mycobacterium *ulcerans* in two isomeric forms (A and B in a 60:40 ratio), only isomer B is cytotoxic (10). Focusing on the primary source of the toxin’s cytotoxic effects, we looked for differences in how the A and B isomers associate with both the Sec61 translocon as well as the ER membrane. The latter was deemed relevant since it likely serves as a reservoir for the toxin prior to association with the translocon. This role is supported by early imaging studies showing uptake into the ER membrane within minutes of the toxin being exposed to cells (11), as well as mycolactone’s strong association with lipophilic carriers, including cellular membranes and HDL (13, 14, 15, 27).

First focusing on the ER membrane, we calculated free energy profiles for toxin association and permeation. Both isomers show a strong association with the ER membrane with energy minima corresponding to configurations that maximize polar and nonpolar interactions at the interface of the lipid headgroups and tails. However, isomer B has a stronger affinity for the membrane than isomer A (∼-17 vs -13 kcal/mol, respectively), with clear differences in the associated conformational ensembles (**Figures 2,3**). The origin of this can be traced to the structure. Mycolactone A has a more collapsed structure, enabling more intramolecular interactions that stabilize the toxin in any environment. In contrast, mycolactone B is necessarily more extended, enabling and requiring more interactions with the environment. Given its amphiphilic nature, this leads to a greater stabilization of mycolactone B compared to A when the toxin transitions from water to the membrane, where both polar and nonpolar interactions are optimized.

The isomeric differences are additionally supported by the average interaction energies, which are stronger between the toxin and membrane for isomer B (more extended), and stronger for intramolecular interactions for isomer A (more compact) (**Table 1**). This also explains the broader angle distribution in the free energy profile of isomer A (**Figure 2**). More intramolecular interactions result in less angular dependence, or need for polar interactions with the lipid head and/or glycerol groups. In contrast, mycolactone B’s extended conformation needs interactions with the lipid headgroups/glycerols to satisfy the polar tails, leading to a narrower angle distribution.

Collectively, isomer B has a stronger water-to-ER membrane association free energy due to its extended conformation, which enables more amphiphilic interactions with lipids. In comparison, isomer A is relatively more stable in water due to its intramolecular interactions. In recent work (15), we also demonstrated that isomer B preferentially associates with the ER membrane containing only POPC and POPE lipids over the more rigid plasma membrane containing POPC, POPE, DPPC and cholesterol. The preferential association was driven by more favorable enthalpic interactions with water in the ER membrane and unfavorable disruption of lipid packing in the plasma membrane. The toxin pulled in the ER-lipids (POPC and POPE) in the plasma membrane, pushing away the saturated DPPC tails and cholesterol. Thus, we hypothesize that isomer B will preferentially localize to more disordered lipophilic carriers within the host, and that this could contribute to its localization to the ER. Our findings here extend this hypothesis to suggest that isomer B will be more likely to localize to the ER, and this increased localization in the membrane containing the Sec61 translocon could contribute to its toxicity via increased local population. An intriguing complication to this hypothesis is that increased affinity for the ER membrane will compete with association with the translocon. In other words, isomer B will have to be ∼3 kcal/mol more stable in the translocon over isomer A, in order to retain the same binding affinity for equivalent ER populations.

Turning to the Sec61 translocon, our simulations reveal clear differences in how each isomer associates with the protein. Mycolactone B was found to have interactions with the plug domain and both sides of the lateral gate of the channel while mycolactone A solely interacts with one side of the lateral gate. Mycolactone A’s lack of interactions with the plug and TMs 7 side of the lateral gate likely reduces its ability to lock these gating elements in place to inhibit translocation. We also find that the B isomer inserts further into the pore constriction of the channel compared to isomer A to form hydrophobic interactions and potentially block the pore from client proteins. Furthermore, key polar interactions of the toxin were found to be better satisfied with mycolactone B bound relative to A, which likely results in a stronger binding affinity of isomer B – making it less susceptible to being displaced by a strong signal sequence. Lastly, the complex bound with the B isomer adopts a relatively more closed conformation on the cytosolic side of the translocon, potentially making it more difficult for an incoming signal sequence to interact with the lateral gate for subsequent translocation.

Based on the simulations in this work there are clear differences in the association of the mycolactone isomers with both the ER membrane and the Sec61 translocon, suggesting that both localization and association could play a role in isomer B’s increased cytotoxicity. Specifically, we find that isomer B has a greater affinity for the disordered ER membrane, which could lead to an increased local concentration around Sec61, increasing its effective affinity for the translocon. Additionally, we show that isomer B’s association with Sec61 better enables the proposed inhibition mechanism in which the inhibitor acts by stitching the key gating elements together at their interface. Finally, we find an induced conformational change of the translocon by mycolactone B that may enhance the isomer’s ability to block the translocation of peptides with a stronger targeting signal. In addition to providing a better understanding of mycolactone’s cytotoxicity, these findings may be informative for the complex mechanism of protein translocation by the Sec61 translocon and the modes of inhibition, as the exact mechanism of many Sec61 inhibitors are unknown. This insight is relevant to targeting specific interactions in the Sec61 translocon for therapeutic applications such as anticancer and antiviral treatment, as cancer cells and viruses rely on and exploit protein translocation into the ER to cause disease (28, 29, 30, 31, 32). Additionally, our findings suggest that diagnostics targeting the mycolactone toxin with disordered lipophilic architectures should have an increased affinity for isomer B. Future experimental work testing these findings by quantifying the relative binding affinities of the two isomers to both the Sec61 translocon and different membrane compositions, as well as the inhibition levels of both isomers will be important additions to the field.

## MATERIALS AND METHODS

The ER membrane was built with 198 units of 1-palmitoyl-2-oleoyl-sn-glycero-3-phosphocholine (POPC) and 102 units of 1-palmitoyl-2-oleoyl-sn-glycero-3-phosphoethanolamine (POPE) using the CHARMM-GUI Membrane Builder for Mixed Bilayers (33). The lipid parameters were obtained from an Amber-based force field (34), with corrections to stabilize the hydrophilic and hydrophobic forces. TIP3P water molecules (35) were added to solvate the membrane with a hydration ratio of 60 molecules per lipid component. This created system boxes with 9.6 nm x 9.6 nm x 9.3 nm (*X, Y, Z*) that were energy minimized for 10,000 steps, heated until 310.15 K, and then equilibrated for 250 ns until reached convergence. After the equilibration, a mycolactone molecule (one isomer per box) was randomly added to the water solution with arbitrary orientation. The toxin parameters (for each isomer) were the same developed and utilized by López et. al. and Aydin et. al. (12, 14). For each isomer, three replicas were created and simulated.

The Newtonian equations of motion were integrated with a time step of 2 fs. During the equilibration and production, the pressure was kept constant at 1 bar (semiisotropically – as recommended for bilayer simulation) by the Berendsen barostat (36) with a coupling time every 5 ps. At the same time, the temperature was controlled at 310.15 K by the velocity rescaling thermostat (7) with a separately coupling time of 1 ps for each component (membrane + toxin and water). Using a cutoff distance of 1.0 nm, the short-range interaction list was updated every 10 steps, along with the long-range electrostatic interactions that were calculated using the smooth particle mesh Ewald method (37). LINCS, the linear constraint solver, was applied to all hydrogen bonds.

The mycolactone-Sec61 complexes were built using Itskanov et al.’s (22) cryo-EM structure of the human Sec61 channel inhibited by mycolactone B. To obtain a complex with the A isomer, AutoGrid v4.2.6 and AutoDock v4.2.6 (38, 39, 40) were used to predict the binding mode of the toxin. AutoGrid was used to set up the toxin and protein and AutoDock was used to perform the docking simulations. Mycolactone B’s position on the protein in Itskanov et al.’s (22) structure was used as a reference to center the grid for docking. The grid was centered on coordinates: 66.249, 55.65, 55.241 Å (*X, Y, Z*) and then expanded to encompass surrounding residues to explore multiple potential binding sites in the region. The grid spacing was set to 0.260 Å, and the grid maps were adjusted to 86 x 92 x 94. Docking was carried out using a Lamarckian Genetic Algorithm (LGA), starting with an initial population of 150 random individuals. The LGA runs included a maximum of 27,000 generations with a maximum number of 2,500,000 energy evaluations. The mutation and crossover rates were set to 0.02 and 0.08, respectively, and an optional elitism parameter of 1 was employed to determine the number of top individuals that would survive into the next generation. The probability of performing a local search on an individual was 0.06 and a maximum of 300 iterations per local search was allowed. The maximum number of consecutive successes or failures before doubling or halving the search step was set to 4. In total, 10 LGA runs were performed. After the conformational search, the docked conformations were sorted based on energy. Finally, the coordinates of the lowest energy conformation were clustered using a root-mean-squared deviation of 2.0 Å. The lowest energy structure of mycolactone A which exhibited a similar binding orientation as mycolactone B in Itskanov et al.’s (22) structure was chosen to be modeled in MD.

Each isomer-Sec61 complex was then embedded in our model ER membrane using CHARMM-GUI Membrane Builder (Bilayer Builder) (33). The POPC and POPE lipids were represented with the Slipids 2020 force field (41, 42) and the Sec61 protein parameters were described by the Amber 99SB-ILDN force field (43). TIP3P water molecules were again used to solvate the complexes. The toxin parameters were obtained from previous work (12, 14, 44). Minimization and equilibration were carried out using a seven-step protocol suggested by CHARMM-GUI for the built system. The systems were energy minimized for 1,000 steps of steepest descent energy minimization followed by 2,000 steps of conjugate gradient energy minimization, using the steepest descent algorithm as the first step in both cases. A six-step equilibration was then carried out, which consisted of gradually removing position restraints on the lipids, protein, and toxin followed by a final equilibration run without any restraints. In all equilibration steps, the system was simulated for 100 ps using the NPT ensemble with temperature and pressure coupling at 310.15 K and 1 bar, respectively. Production was performed under the same conditions as described above.

All MD simulations – toxin with ER membrane or complexed with Sec61 – were performed with the GROMACS 2019.4 package (45). In-group developed *Python* scripts were used to calculate the hydration and obtain the two-dimensional Potential of Mean Force maps (2D-PMFs) for the membrane permeation process. MDAnalysis (46, 47) was also used in Python scripts to perform contact analysis for the mycolactone-Sec61 complexes. VMD (48) was used to visualize the trajectories and generate the Figs. Finally, the GROMACS was patched with PLUMED 2.5.3 (49) to enable the employment of the enhanced method TTMetaD.

Membrane permeation is generally a slow and challenging process to be captured by MD simulations on accessible timescales. Enhanced samplings methods, such as Metadynamics (50, 51), were developed to address this and allow the observation of this type of process. TTMetaD is an efficient variant of Metadynamics that enables the convergence of the system with only an approximated idea of the barrier surroundings, not its height (52). This information can be obtained by unbiased simulations of the toxin permeation, e.g., in our previous studies (14, 15). In a simulation with MetaD, a bias energy is added to the Hamiltonian of the system through a small number of selected CVs that represent the slowest degrees of motion of the system. This bias energy is usually a Gaussian function that is centered at the previous visited configuration and represents the amount of energy necessary to sample the transition state in the CV space. In TTMetaD, the bias energy incremental rate decreases exponentially with respect to the local bias, first filling the transition state region, and only then converge through a more aggressive tempering of the Gaussian height (52, 53). Based on previous studies of small organic molecules (53, 54) and specifically for mycolactone (14, 15), two CVs are enough to describe the permeation process: the *z*-component distance of the center of mass of the lactone ring to the COM of the membrane as CV1; and the angle between the vector connecting the northern and southern chains’ hydroxyl groups and the normal vector to the bilayer. More detail can be seen in references (14, 15).

As an efficient method for membrane permeation analysis (14, 53, 54), 2D PMFs were obtained to observe the energy distribution along the permeation process and to allow the calculation of the minimal free energy path (MFEP). The 2D-PMF is computed by the reverse of the average bias energy from the independent replicas followed by a diagonal symmetrization with respect to the CVs (toxin orientation and the center of the membrane). The MFEP was calculated using the zero-temperature string method (55) and represents the most probable path in the ensemble of permeation processes. One-dimensional (1D) free energy profiles were obtained by taking the average MFEP from the simulations for each isomer.

## SUPPORTING MATERIAL

Supporting Material can be found online at https://doi.org/XX.YYYY/j.bpj.2023.0Z.ZZZ

## AUTHOR CONTRIBUTIONS

J.M.J.S., G.C.A.H and J.D.M.N designed the research. G.C.A.H and J.D.M.N performed the simulations and analyses. All authors interpreted the results and wrote the manuscript.

## DECLARATION OF INTEREST

The authors declare no competing interests.

## ACKNOWLEDGEMENTS

The authors thank Professor Eunyong Park for kindly providing the coordinates to their sec61 structure prior to final publication and for helpful discussions. We gratefully acknowledge support from the National Institute of General Medicine of the National Institutes of Health under award number R35GM143117 and computational support from the Extreme Science and Engineering Discovery Environment supported by the National Science Foundation (Grant No. ACI-1548562) under allocation MCB200018 as well as the Center for High Performance Computing at the University of Utah.

## Notes

### Competing Interest Statement

The authors have declared no competing interest.

