## Supplemental Figures 1-3 for "Mycolactone A vs. B: Does localization or association explain isomer-specific toxicity?"

### SUPPORTING INFORMATION

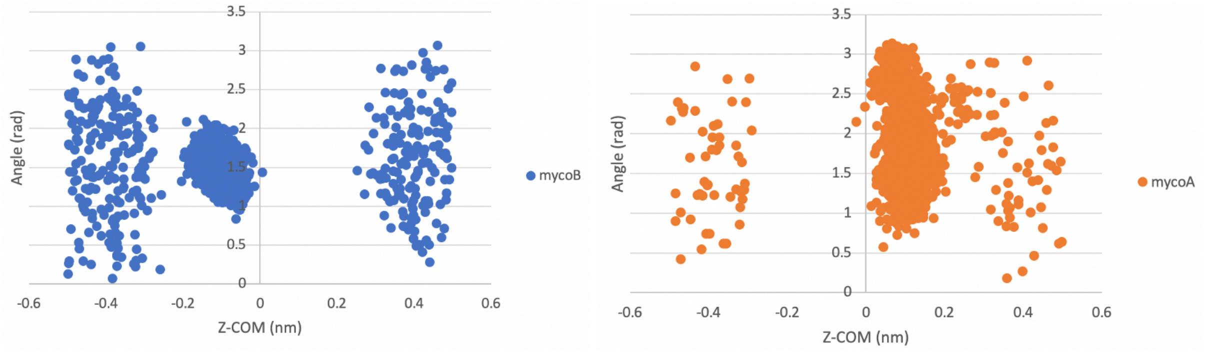


**Figure S1**. Distribution of the two CVs in unbiased simulations of mycolactone A/B with the ER membrane.

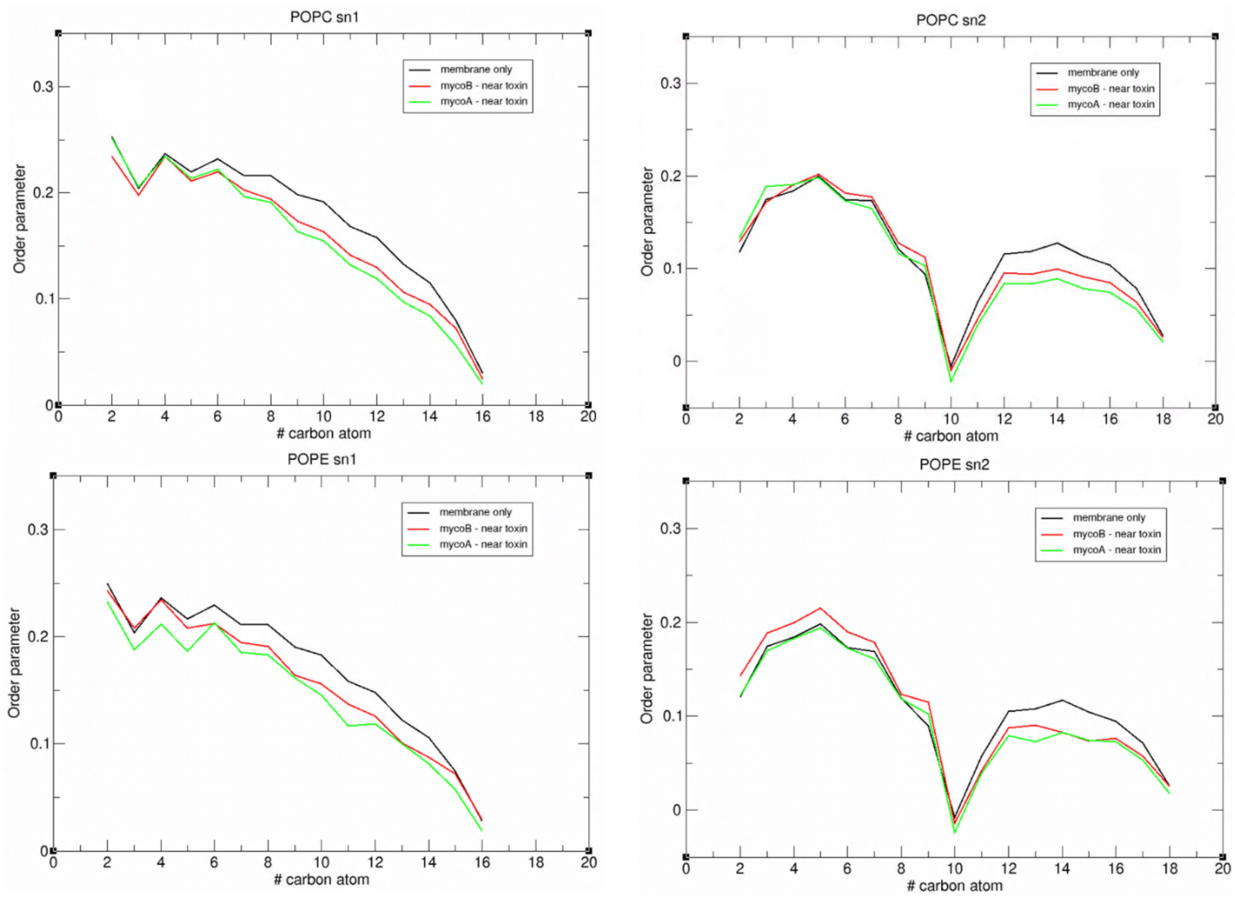


**Figure S2**. Tail order parameters of lipids near mycolactone and lipids of an ER membrane without the toxin.

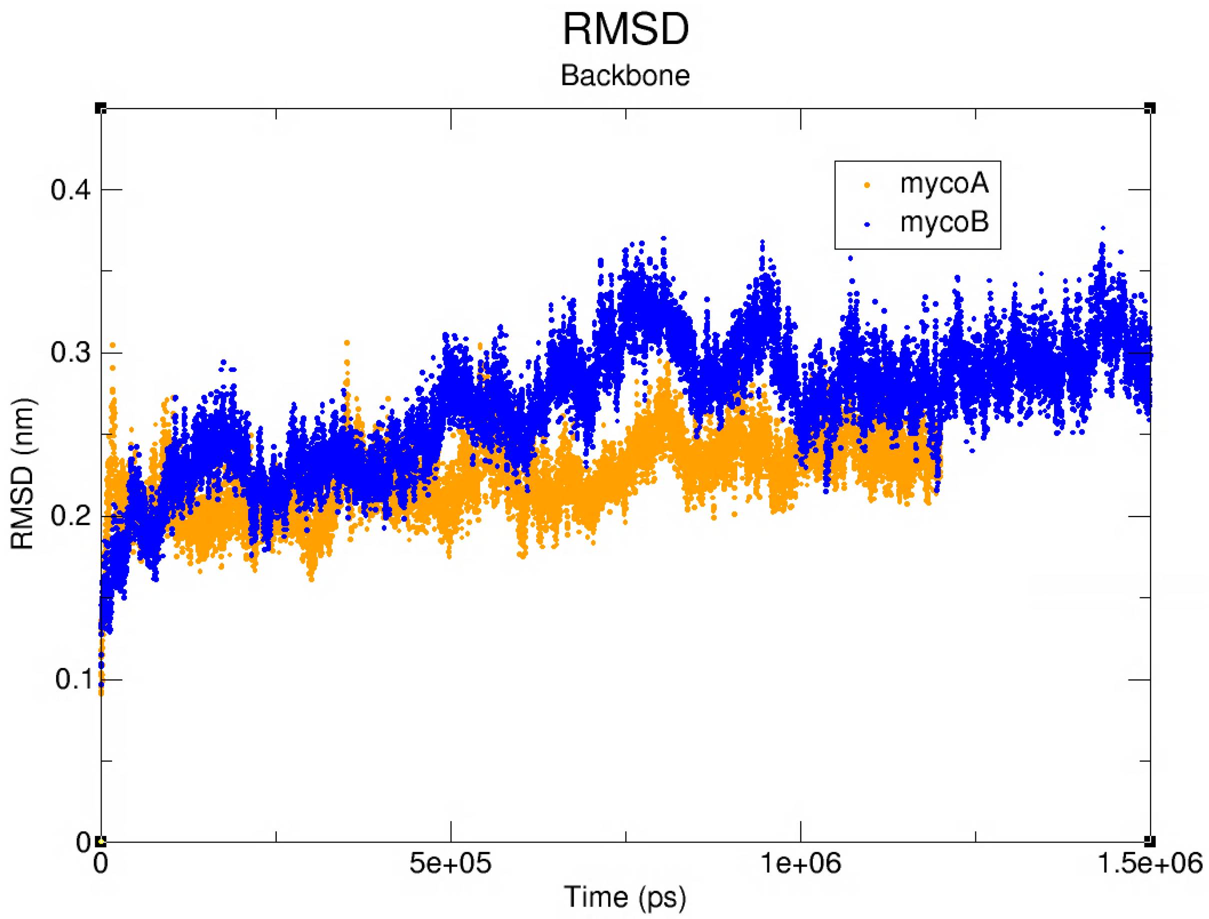


**Figure S3**. Root mean square deviation of mycolactone-Sec61 complexes.
